# Three Subtypes of Autism Spectrum Disorder with Transcriptomic Signatures Derived from Morphometric Similarity Networks

**DOI:** 10.1101/2024.09.06.611561

**Authors:** Hongxiu Jiang, Raul Rodriguez-Cruces, Ke Xie, Valeria Kebets, Yezhou Wang, Clara F. Weber, Ying He, Jonah Kember, Jean-Baptiste Poline, Danilo Bzdok, Seok-Jun Hong, Boris Bernhardt, Xiaoqian Chai

## Abstract

Autism spectrum disorder (ASD) is a prevalent and highly heterogeneous neurodevelopmental disorder. Previous studies have attempted to identify ASD subgroups by analyzing isolated cortical structural features. However, these studies have not considered the relationship between multiple structural features, which provide information on the structural organization of the brain. Morphometric similarity network (MSN), a structural brain network contributed by multiple anatomical features (gray matter volume, mean cortical thickness, surface area, mean curvature, Gaussian curvature, curvature index, and fold index), strongly relates to cytoarchitectonic and genomic measures of histological similarity between cortical areas. We applied K-means clustering on MSN from 236 individuals with ASD and identified three subtypes. Subtype-1 showed relatively similar MSN values with typically developmental individuals (TD). Subtype-2 showed higher morphometric similarities in the lateral frontal and temporal cortical regions and lower in anterior prefrontal and occipital regions compared to TD. These patterns were the opposite in subtype-3. Behaviorally, subtype-3 had more severe verbal and social deficits compared to subtype-2. The weaker resting-state functional connectivity (rs-FC) between the language and salience networks was observed between subtype-2 and TD. Subtype-3 had stronger rs-FC between salience and default mode networks (DMN), between frontoparietal and visual networks, and between language and dorsal attention networks, while weaker rs-FC within DMN, within sensorimotor, and within salience networks. In addition, genes with expression patterns associated with regional MS changes in ASD subtypes were functionally enriched in neuron-specific biological processes related to nervous system development, synaptic signaling and chromatin organization. These genes were particularly enriched in GABAergic neurons, glutamatergic neurons, astrocytes and microglia. Taken together, our findings suggest the existence of different neuroanatomical subtypes based on multiple anatomical features in ASD with distinct transcriptomic signatures and functional connectome patterns.

## INTRODUCTION

Autism spectrum disorder (ASD) is a prevalent developmental disorder, characterized by impairments in social communication, restricted interests, and repetitive behaviors. The current ASD diagnostic criteria use a series of behavioral measurements to identify individuals with autistic traits. However, among ASD individuals, there is great variability in both behavioral patterns and underlying biological causes (London, 2014). Previous studies have investigated atypical brain structural and functional development using magnetic resonance imaging (MRI) but findings from these studies have been inconsistent (Chen et al., 2011; Hyde et al., 2010; Khundrakpam et al., 2017; Pua et al., 2017). High heterogeneity within ASD is a major contributing factor to this inconsistency, in addition to methodological variation in sampling and image acquisition and analyses. To overcome the challenge of heterogeneity, researchers use data-driven strategies to identify subgroups of ASD with shared common structural and/or functional patterns. For example, Chen et al. (2019) identified three subtypes by voxel-based morphometry conducted by gray matter volume maps. Zabihi et al. (2020) reported five subtypes of ASD by normative modeling of cortical thickness. In addition to these works based on single anatomical features, Hong et al. (2018) found three distinctive anatomical subtypes based on cortical thickness, intensity contrast, cortical surface area and geodesic distance. These voxel- or vertex-based methods provided preliminary evidence for brain-based subtypes in ASD.

A relatively consistent finding in brain imaging study of ASD is the atypicality of large-scale structural networks (Bernhardt et al., 2016; Jiang et al., 2023). Compared to more traditional structural networks constructed based on structural covariance and tractography, morphometric similarity networks (MSN) proposed by Seidlitz et al. (2018), have higher correspondence cortical cytoarchitecture and can be constructed for individual subjects. Morphometric similarity, which measures the inter-regional morphometric similarity of multiple morphometric features, strongly relates to gene co-expression between cortical areas (Seidlitz et al., 2018). MSN has led to new insights into different psychiatric disorders. For instance, (Morgan et al., 2019) found that genes that were up-regulated in post-mortem brain tissue from patients with schizophrenia are normally overexpressed in association cortical areas that have reduced morphometric similarity in psychosis. (Li et al., 2021) reported the expression of MDD-associated genes spatially correlates with MSN differences.

Here, we identify subtypes in ASD based on MSN, then link MSN abnormalities of each subtype to transcriptomic data and functional connectivity patterns to advance our understanding of the multimodal cortical organization and molecular underpinning of heterogeneity in ASD. First, we constructed MSN matrices and used K-means clustering to obtain three subtypes. Second, we identified differences between each subtype and typically developing individuals (TD), and between subtypes on MSN patterns, behavioral scores, and resting-state functional connectivity patterns. Third, we applied partial least square regression (PLSR (Wold, 1966)) to explore the relationship between brain-based differences and genetic signatures from the Allen Human Brain Atlas of each subtype and performed a genetic functional enrichment analysis and a cell-type enrichment. Our results support the existence of different neuroanatomical subtypes in ASD with distinct genetic signatures.

## METHODS

### Subjects

Subjects from the Autism Brain Imaging Data Exchange (ABIDE-1) dataset (Di Martino et al., 2014) and ABIDE-2 (Di Martino et al., 2017) were selected using the following inclusion criteria: 1) male; 2) subjects with Autism Diagnostic Interview-Revised (ADI-R) scores, or the Autism Diagnostic Observation Schedule (ADOSO), or the Autism Diagnostic Observation Schedule Second Edition (ADOS-2); 3) acceptable structural MRI quality: T1-weighted (T1w) images were visually inspected for motion artifacts, and the Euler Number (Dale et al., 1999; Monereo-Sánchez et al., 2021; Rosen et al., 2018) of the reconstruction images by FreeSurfer is 2; 4) acceptable functional MRI quality (details in *MRI preprocessing and Functional Connectivity Construction*).

A total of 236 individuals with ASD and 287 typically developing (TD) individuals were included in the data analysis. Across sites, ASD and TD had similar ages (ASD: mean ± SD = 16.61 ± 9.01, range = 5.1 – 57; TD: mean ± SD = 16.61±10.17, range = 5.9 – 62; t = -0.004, p = 0.996). Compared to TD, individuals with ASD had lower full-scale IQ (ASD: 105.54±16.42, range = 65 - 148; TD: 113.9±12.03, range = 79 - 148; t = -6.97, p < 0.001).

### MRI preprocessing and Functional Connectivity Construction

All sites provided T1-weighted (T1w) MRI and resting-state functional MRI (rs-fMRI) that were scanned using 3T Siemens (New York University Langone Medical Center (NYU), Oregon Health and Science University (OHSU), University of Pittsburgh School of Medicine (PITT), University of Utah School of Medicine (USM)), Philips (Barrow Neurological Institute (BNI), Kennedy Krieger Institute (KKI)) or General Electric (San Diego State University (SDSU)) scanners (see Supplementary for details).

T1w MRI images were processed using FreeSurfer (v6.0, https://surfer.nmr.mgh.harvard.edu/). Skull-stripping, bias field correction, registration, anatomical segmentation, and surface reconstruction were applied to structural images. The cortical surfaces were divided into 308 regions (Romero-Garcia et al., 2012), which were derived from the 68 cortical regions in the Desikan-Killiany atlas. For each region, seven morphometric features were employed in the MSN construction (see below for details): gray matter volume (GM), mean cortical thickness (CT), surface area (SA), mean curvature (MC), Gaussian curvature (GC), curvature index (CI), and fold index (FI) (Li et al., 2017).

rs-fMRI data were processed and analyzed using the CONN-fMRI Functional Connectivity toolbox v20b (Whitfield-Gabrieli & Nieto-Castanon, 2012), MATLAB R2021b. All sequences were preprocessed using the default pipeline of the CONN toolbox: functional label current files as “original data”, functional realignment and unwarp, functional center to (0,0,0) coordinates, functional outlier detection (Artifact Detection Tools was used to identify the outlier scans for scrubbing), structural center to (0,0,0) coordinates, functional indirect segmentation and normalization, functional label current files as “MNI-space data”, and functional smoothing (spatial 8-mm Gaussian kernel). Data were denoised, and signal data from the cerebrospinal fluid and white matter were regressed and processed with a bandpass filter (0.008–0.09 Hz) to reduce the noise effects from physiological effects and scanner drift. Subsequently, the preprocessed data was subjected to ROI-to-ROI connectivity analyses and connectome matrices were derived to visualize the results. We selected two sets of ROIs to verify the differences among participants: 1) seven network-level ROIs defined by (Yeo et al., 2011); 2) 32 ROIs defined from CONN’s ICA analyses of the HCP dataset. By extracting the average mean time series of ROIs, the correlation coefficient between the mean time series of any two ROIs was calculated and converted using Fisher’s r-to-z transformation.

### Morphometric Similarity Network (MSN) Construction

In each participant, the MSN was constructed by calculating Pearson’s correlation for each pair of z-normalized morphometric feature vectors consisting of the seven morphometric features of different regions, forming a 308*308 matrix for each participant (Li et al., 2021; Seidlitz et al., 2018), as shown in **Fig. 1A**. Then, the mean regional MSN was calculated by averaging the values in each row and formed a vector of length 308 for each subject. We implemented NeuroCombat (Fortin et al., 2018) to adjust for site effects. ComBat corrects potential scanner/site effects on the brain data by harmonizing the mean and variance of brain measures across scanners. Then we computed a Linear Regression Model (LRM) to account for the age effect, and used the residuals as the corrected mean regional MSN values for subsequent analyses.

**Fig. 1.**
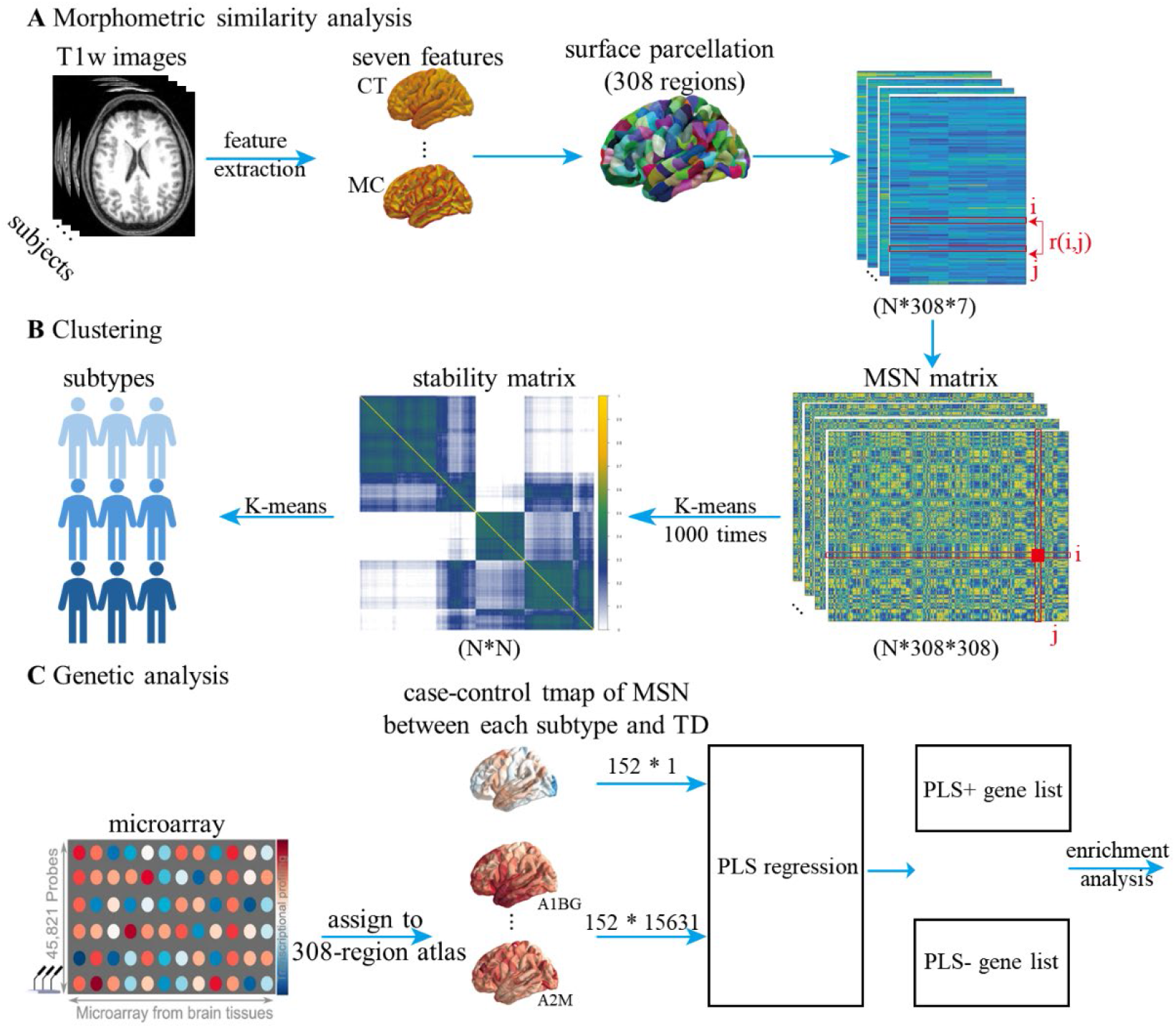
Methods overview. **A** Structural features extraction and morphometric similarity networks construction. **B** The clustering process. **C** Genetic analysis.

### Clustering Analysis

We used the residual MSN features from the LRM to run the K-means (MacQueen, 1967) for clustering, with the number of clusters *k* ranging from 2 to 10. To obtain stable subtyping results, we carried out 1,000 iterations of bootstrapping and constructed the stability matrices (Rodríguez-Cruces et al., 2020) for each cluster number. In each bootstrapping iteration, a binary N*N stability matrix would be generated (N is the number of subjects). If two subjects from a specific row and a column are classified into the same cluster, one is placed in that cell. A zero in the cell means the subjects were classified into different clusters. The final stability matrix is the mean matrix of 1,000 matrices, which summarizes the probability that each pair of subjects belongs to the same cluster after 1,000 times of clustering. This bootstrap-based clustering method identifies stable subclasses from the whole ASD group that show a similar MSN characteristic. As shown in Fig. S1, the stability matrices converged to form three clusters. Finally, the stability matrix of k=4 was chosen as the final stability matrix and K-means clustering was performed over the final stability matrix to obtain the ASD subtypes, as shown in **Fig. 1B**.

In addition, principal component analysis (PCA) (Wold et al., 1987) was used on MSN matrices of the whole subjects to visualize. PCA helps identify the key components that capture the greatest amount of variance in the data and then projects the original data onto these components.

### Group comparison analyses (symptom scores and functional connectivity)

To compare MSN maps between each subtype and TD, we ran two-sample t-tests on the corrected mean regional MSN for each region. Behavioral differences were measured by using a one-way analysis of covariance (ANOVA) implemented with the *aov* R function, with a significance level of p < 0.05. A post hoc Bonferroni-adjusted pairwise t-test was used to compare the scales of the specific two subtypes.

To understand functional connectivity patterns in each subtype, we performed ROI-based analyses on all subjects with a general linear model to determine significant connections at the individual level (first-level analysis) in CONN. Based on the results obtained in the first-level analysis, a two-sample t-test was performed (controlling for age and site) with a threshold set at p < .05 false discovery rate (FDR) corrected, to determine significantly different connections between TD and ASD subtypes (second-level analysis).

### Regional gene expression matrix

We used transcriptomic data from the Allen Human Brain Atlas (AHBA, http://human.brain-map.org/; (Hawrylycz et al., 2012)), which includes a whole-genome expression atlas from 6 postmortem donors (age = 42.50 ±13.38; male/female = 5/1). We used the *Abagen* toolbox (Markello et al., 2021) to preprocess the transcriptomic data with the default setting. In brief, genetic probes were reannotated using updated information provided by Arnatkevičiūtė et al. (2019) to remove probes with less reliability rather than the default one provided by from AHBA dataset. The reannotated probes were kept only when their intensity related to the background noise level was larger than 50%. Then microarray expression data was normalized and aggregated (Markello et al., 2022) to the above-mentioned 308-region parcellation (Romero-Garcia et al., 2012). Because only two right hemisphere data are available, this study analyzed the left hemisphere with 152 regions. These steps result in the gene expression matrix with the dimension of 152 * 15, 631 for the analysis.

### Linking MSN to gene expression in ASD subtypes

We applied *plsregress* in MATLAB to run PLS regression (PLSR) analysis, linking the difference of MSN between each subtype of ASD and TD, to gene expression data, **Fig. 1C**. The gene expression data were used as the predictor variable (i.e., 15,631 input variables) across the 152 region observations, while the t-statistic map between ASD subtype and TD in 152 left regions was used as the response variable (i.e., one outcome variable). In brief,) the PLSR analysis is to decompose the predictor (i.e., 15,631 input variables (genetic expression matrix) to create orthogonal components that can best predict) across the outcome variables (MSN tmap differences).152 region observations We only focus on the first component of PLSR (PLSR1), as it explains the majority of the variance. A permutation test was further used to test the statistical significance of the variance explained by PLSR1, where the spatial permutation models generate spatial-constrained null distribution by applying random rotation to spherical projections of the brain (Alexander-Bloch et al., 2018). Bootstrapping was used to estimate the variability of each gene’s PLSR1, with the Z score of the ratio of the weight of each gene to the standard error of bootstrap as the criteria of ranking their contributions to PLSR1.

### Enrichment Analysis of Genes Transcriptionally Related to Regional Changes

The obtained PLSR1 included PLSR1+ (the greatest positive weight) and PLSR1-(the greatest negative weight), and the corresponding gene lists were inputted to functional enrichment analysis including gene ontology (GO) biological processes, Kyoto Encyclopedia of Genes and Genomes (KEGG) pathways, Reactome Genes Sets and WikiPathways on the gene lists using Metascape (Zhou et al., 2019). The expression-weighted cell type enrichment (Skene & Grant, 2016) was carried out to identify the differences between two subtypes in cell type, with the PLSR1 gene lists as inputs.

## RESULTS

### Altered MSN in three ASD subtypes

We identified three subtypes of ASD based on MSN values. **Fig. 2A** shows the distribution of the first two principal component analysis (PCA) components of MSN in TD and three ASD subtypes. In both the scatter plot and smoothed density histogram, TD individuals had the largest range of PCA1 values. Subtype-2 was characterized by positive PCA1 values, while subtype-3 was characterized by negative values. Subtype-1 has the narrowest range of PCA1 values that were slightly negative.

**Fig. 2.**
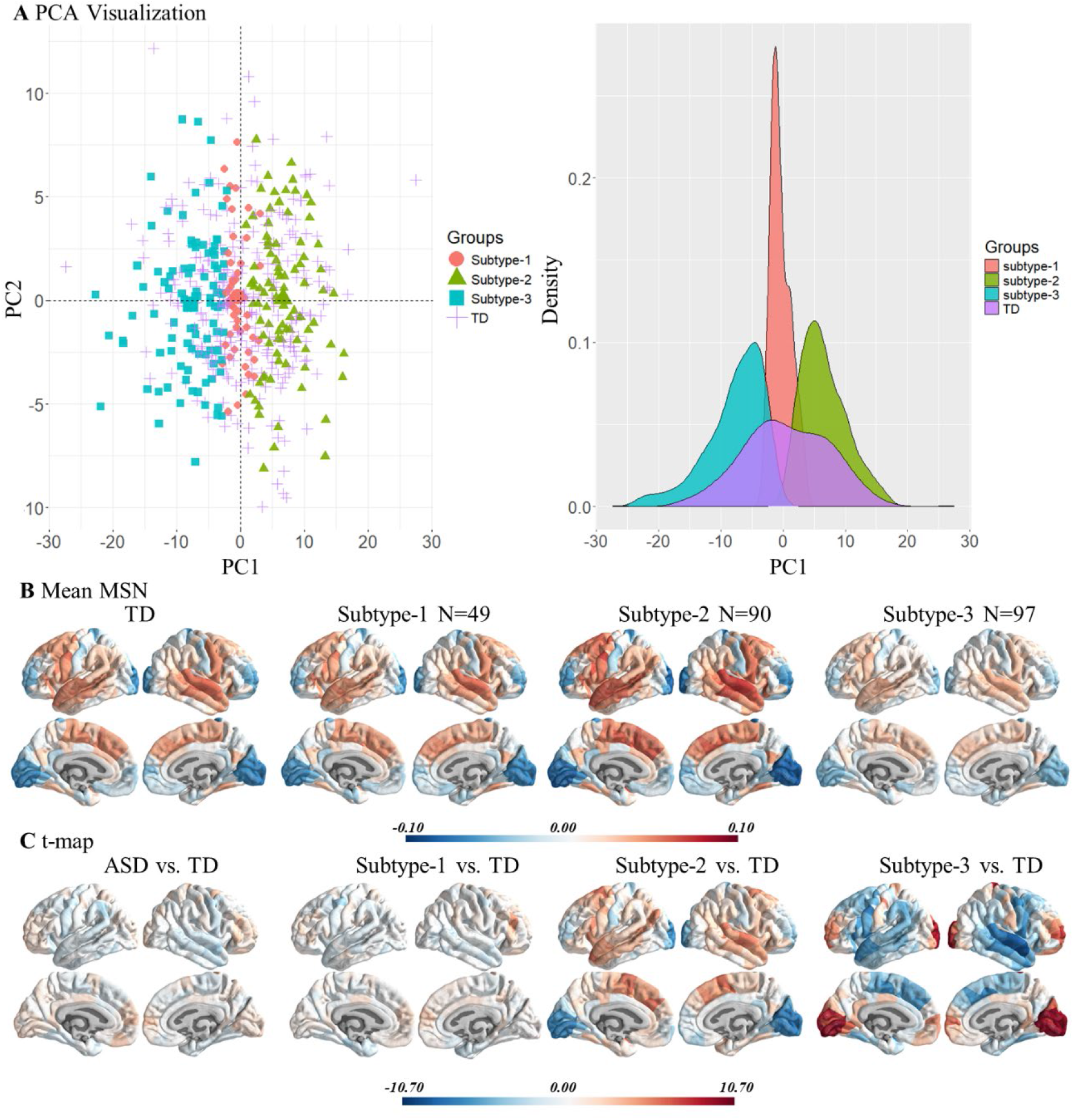
The visualization of individuals and the case-control differences in mean regional morphometric similarity networks (MSN). **A** The scatter plot of TD and subtypes of ASD, the X and Y axes are the first two principal components of the mean regional MSN matrix of each group of participants (the left); the distribution curve of the first principal component of the MSN matrix (the right). **B** The mean regional MSN of each group (corrected for age and site). **C** Corrected case-control The t-map of the mean regional MSN.

The mean regional MSN of the TD group and the three subtypes of ASD are shown in **Fig. 2B**. Higher regional MSN implies higher morphometric similarity between an area and the rest of the cortex, and conversely for lower regional MSN. **2B**.

The pattern results of group-level mean MSN maps of the TD group were similar to previous studies (Li et al., 2021; Morgan et al., 2019). Specifically, the. The caudal middle frontal, superior frontal, superior temporal and middle temporal cortices showed positive MSN values, and lateral occipital and lingual cortices showed negative MSN values in TDs. Higher regional MSN implies higher morphometric similarity between an area and the rest of the cortex, and conversely for lower regional MSN.

The mean regional MSN of the ASD subtype-1 was similar to TD, while subtype-2 and subtype-3 were different from TD in the above-mentioned regions. Compared to TD, subtype-2 had more positive values in regions where the MSN values of TD were positive (e.g. temporal and frontal regions), and more negative values in regions where the TD values were negative (e.g. occipital cortex). Subtype-3 had an oppositive pattern in these regions, specifically, less positive MSN values in regions with positive values in TDs and less negative values in regions with negative values in TD. Two sample t-tests between individuals in each subtype and TD group (**Fig. 2C**) highlighted these differences.

### Clinical symptom severity across three ASD subtypes

Using an ANOVA, we found significant differences in the ADIR-Social (*F* = =3.94, *p* = =0.049) and ADIR-Verbal (*F* = =5.32, *p* = =0.022) in the three ASD subtypes. We then performed post-hoc t-tests to better understand these group differences. The two-sample t-test of symptom severity profiles revealed that subtype-3 had higher scores in ADIR-Social (*t* = -2.49, *p* = 0.04) and ADIR-Verbal (*t* = -2.77, *p* = 0.015) than subtype-2. That is, individuals in subtype-3 have fewer social and verbal deficits according to ADIR scales than subtype-2.

**Table 2.** Symptom severity profiles of subtypes.

|  | ADIR-Social | ADIR-Verbal | ADIR-RRB |
| --- | --- | --- | --- |
| subtype-1 vs. subtype-2 | 0.67 (0.44) | 0.67 (0.46) | 0.29 (-1.18) |
| subtype-1 vs. subtype-3 | 0.13 (-1.70) | 0.07 (-2.14) | 0.11 (-2.33) |
| subtype-2 vs. subtype-3 | **0.04 (-2.49)** | **0.015 (-2.77)** | 0.29 (-1.19) |

Note: p values (t values), after Benjamini & Hochberg correction.

### Atypical functional connectivity patterns of three ASD subtypes

To further explore the distinctions among subtypes, we compared functional connectivity between the subtypes. We calculated the network-wise connectivity using Yeo seven-network atlas. Compared to TD, subtype-1 had significantly stronger functional connectivity (hyper-connectivity) between Salience/Ventral attention and Somatomotor networks (**Fig. 3A)**; Subtype-3, compared to TD, had stronger connections between Salience/Ventral attention and somatomotor networks, Control and Visual networks, Control and Somatomotor networks, Control and Default mode networks. Subtype-2 did not show any significant differences compared to TD. In addition, the results based on network-wise ROIs in CONN showed that subtype-2 had significantly weaker FC (hypo-connectivity) between Language and Salience networks; subtype-3 had stronger FC between Salience and Default mode networks, Frontoparietal and Visual networks, Language and Dorsal attention networks, as well as weaker FC within Default mode networks, within Sensorimotor networks, and within Salience networks, compared to TD (**Fig. 3B)**.

**Fig. 3.**
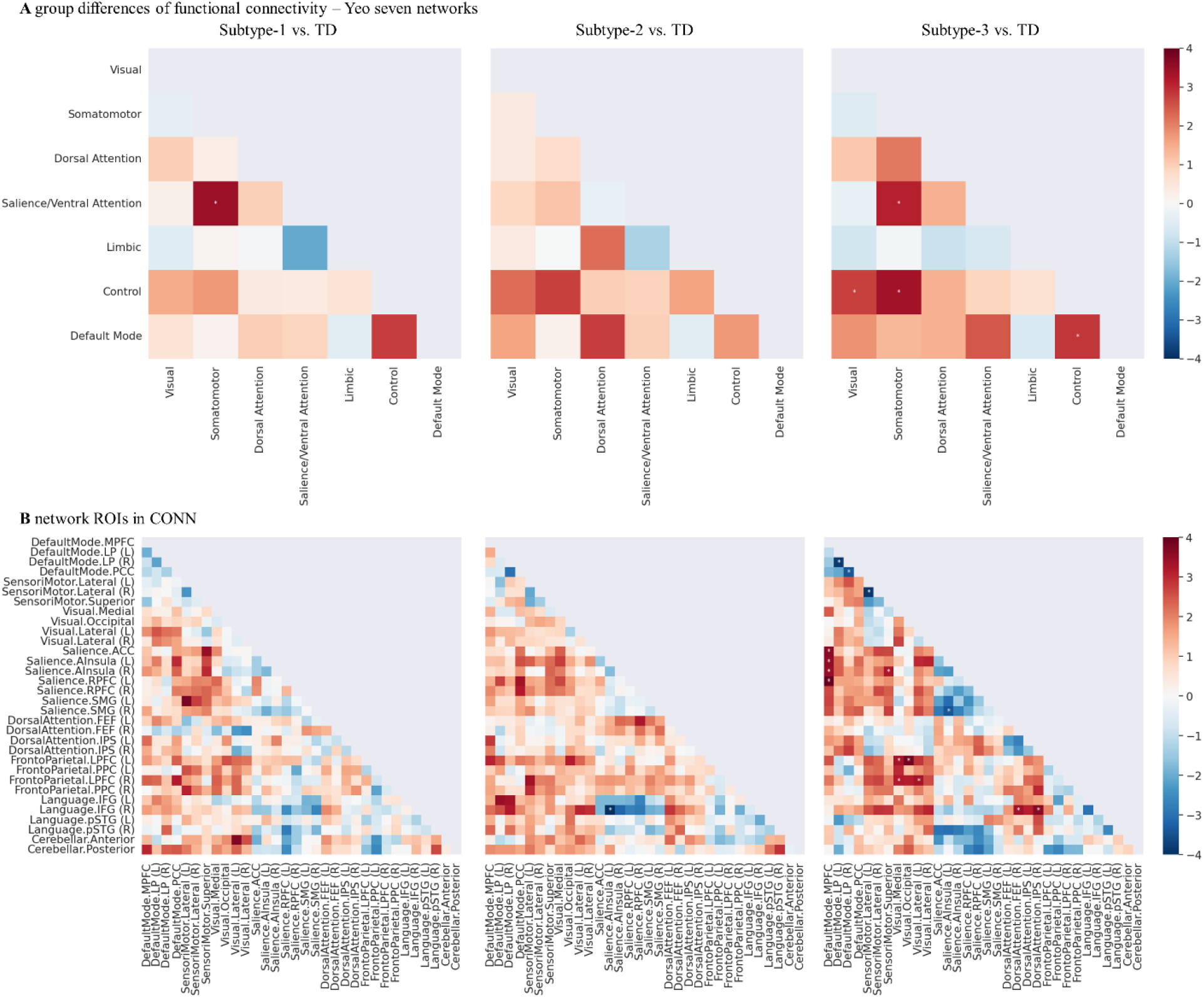
Functional connectivity profile of subtypes. The group differences (t values) of functional connectivity **A** in seven networks, **B** in 32 network ROIs. * represents p<0.05 (false discovery rate (FDR) corrected.)

### Transcriptional profile related to the change of MSN

The genetic data from AHBA was preprocessed by the Abagen toolbox forming the 152 * 15,631 cortical gene expression matrix. Then we used PLSR to explore differences between the differences of MSN and the gene expression matrix among the subtypes. The first component (PLSR1) was defined as the spatial map that captured the greatest fraction of total gene expression variance across cortical areas. PLSR1 of three subtypes explained 12.3%, 47.4%, and 46.6% of the variance, respectively. PLSR1 weighted gene expression map was significantly spatially correlated with the case-control t-map with the Pearson correlation efficient (r=0.35, r=0.69, and r=0.68 respectively) (**Fig. 4A**). These positive correlations suggest that genes positively weighted on PLSR1 are overexpressed in regions where morphometric similarity was increased in individuals with ASD compared to TD, while negatively weighted genes are overexpressed in regions where morphometric similarity was decreased in individuals with ASD compared to TD.

**Fig. 4.**
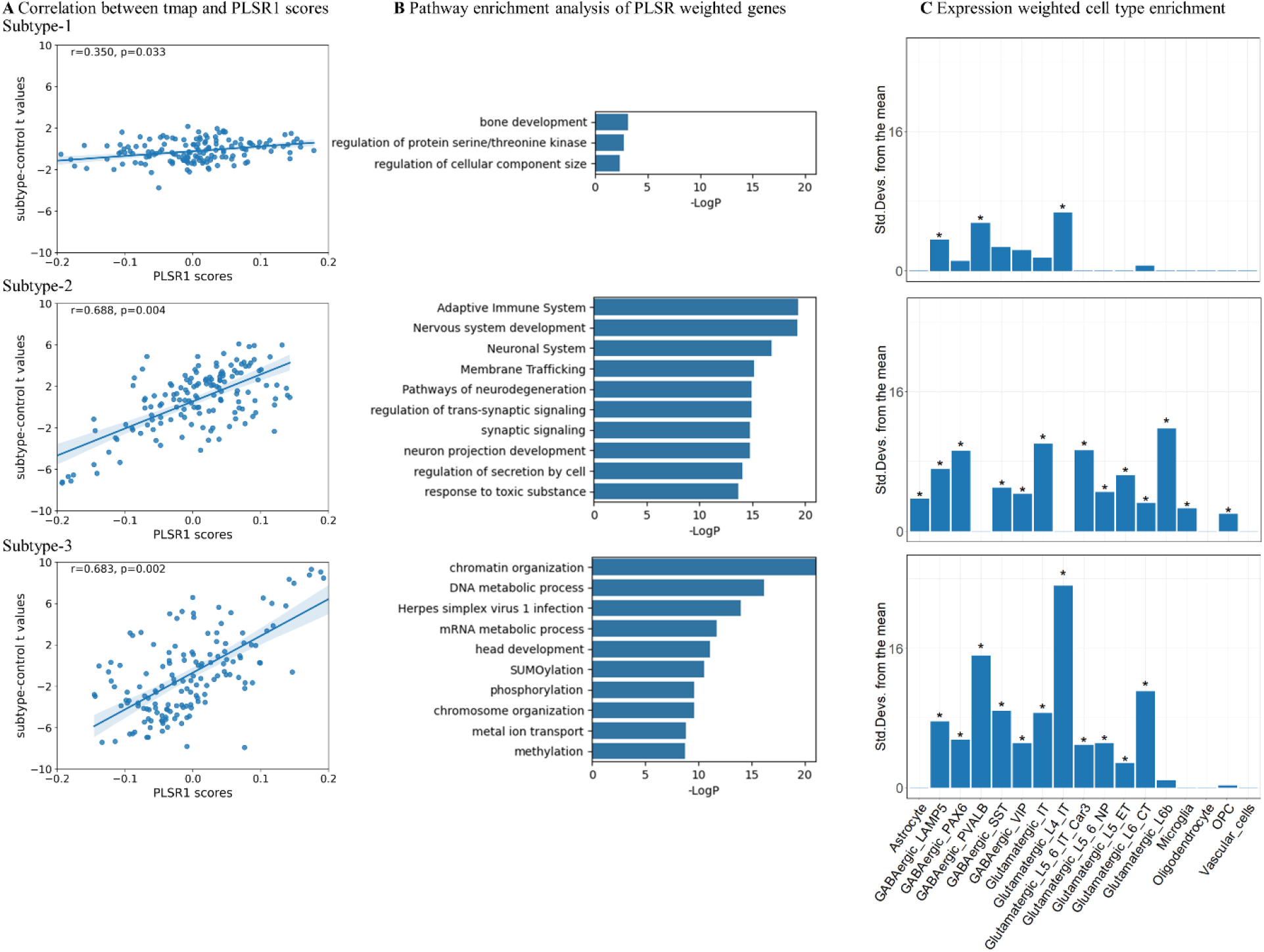
Genetic analysis results. **A**. The correlation between the case-control t-map of regional MSN and PLSR1 scores in each subtype. **B**. The top-20 Gene Ontology biological processes. **C**. The Expression Weighted Cell Type Enrichment. GABA: gamma-aminobutyric acid, LAMP5: Lysosome-associated membrane glycoprotein 5, PAX6: Paired Box 6, PVALB: parvalbumin, SST: somatostatin, VIP: vasoactive intestinal peptide, L4: layer 4, IT intratelencephalic-projecting, CAR3: Carbonic Anhydrase 3 expressing, NP: near-projecting, ET extratelencephalic-projecting, CT corticothalamic, OPC oligodendrocyte progenitor cells.

We ranked the normalized weights of PLSR1 based on univariate one-sample Z tests and found 27 genes (Z>5) of subtype-1, 2102 genes of subtype-2 and 2196 genes of subtype-3. **Fig. 4B** showed the top twenty significant genetic enrichment pathways and processes. Genes for subtype-1 were mostly related to the regulation of protein serine/threonine kinase activity and regulation of cellular component size; genes for subtype-2 were mostly involved in nervous system development and membrane trafficking and synaptic signaling; genes for subtype-3 were mostly related to the processes of chromatin organization, DNA metabolic process and head development. **Fig. 4C** showed cell-type enrichment analysis results: there were two significant subclasses of inhibitory GABAergic neurons (LAMP5 and PVALB), and excitatory Glutamatergic L4_IT subclass neurons for subtype-1; astrocyte, four GABAergic neurons (LAMP5, PAX6, SST and VIP) and six Glutamatergic neurons (IT, L5_6_IT_Cars3, L5_6_NP, L5_ET, L6_CT and L6b), microglia, and oligodendrocyte progenitor cells for subtype-2; and five GABAergic neurons (LAMP5, PAX6, PVALB, SST and VIP) and six Glutamatergic neurons (IT, L4_IT, L5_6_IT_Cars3, L5_6_NP, L5_ET and L6_CT) for subtype-3. **Table S1** provides a brief function description of these neurons.

## Discussion

High heterogeneity is inherent in ASD. Better understanding brain-based subtypes within ASD is a key step toward uncovering the mechanism of this disorder. In this study, we identified three subtypes of ASD and each of them showed a unique pattern in the structure and function of the brain, behavior, and genes. Previous studies mostly used one structural feature (Chen et al., 2019; Liu et al., 2022) or used more than one structural feature without considering the correlation between features (Hong et al., 2018). To our knowledge, this study is the first attempt to use structural networks constructed from multiple structural features to uncover subtypes of ASD.

The regions with MSN differences among subtypes included the caudal middle frontal cortex, superior frontal cortex, superior temporal cortex, middle temporal cortex, lateral occipital cortex, and lingual cortex, which relate to many cognitive functions. The middle and superior frontal cortices are part of the prefrontal cortices and are critical for executive functions including working memory, rule learning and planning (Badre & D’esposito, 2009; Harika-Germaneau et al., 2022). The superior and middle temporal cortices have been characterized by language and multisensory processing (Fan et al., 2017; Petrides, 2023; Stevenson et al., 2016). The lateral occipital cortex is related to object recognition, facial recognition, and motion perception (Palejwala et al., 2020). Differences in these regions have been reported in individuals with ASD compared to typically developing people, including anatomical (Bigler et al., 2007; Chandran et al., 2021) and functional analysis (Delmonte et al., 2013; Jao Keehn et al., 2017; Jung et al., 2019; Xu et al., 2020).

Similarly, the behavioral comparison implicated language- and social-related differences. Functional analysis revealed atypical connections between the salience and default mode networks, salience and language networks, frontoparietal and visual networks, and within default mode networks. These findings are compatible with the idea that ASD is characterized by abnormalities of distributed functional networks, rather than focal impairment (Wang et al., 2019). For instance, the default mode network contributes to processes that include mentalizing, theory of mind, and self-referential processing and plays an important role in the symptomatology of ASD (Padmanabhan et al., 2017). Reduced salience network integrity has been associated with sensory and socio-communicative symptoms (Abbott et al., 2016). Individuals with ASD have higher medial prefrontal cortex connectivity with the insula, a core region of the salience network, and this connectivity negatively predicts the Social Responsiveness Scale (Chen et al., 2022). Individuals with ASD show a reduced ability to integrate visual or multi-sensory feedback during motor behavior (Lepping et al., 2022), and are more reliant on visualization to support language comprehension (Kana et al., 2006).

The transcriptomic analysis revealed genetic differences related to the structural atypicality of the subtypes. Subtype-1 is enriched in cell type GABAergic LAMP5 and PVALB, and Glutamatergic L4_IT. LAMP5 plays a pivotal role in sensorimotor processing in the brainstem and spinal cord in mice models (Koebis et al., 2019), and LAMP5 mutant mice showed decreased anxiety (Tiveron et al., 2016). The GABAergic_PVALB (Parvalbumin) neurons affect the neuronal functions and dynamics by adjusting buffering calcium (Filice et al., 2016; Lawrence et al., 2010). The evidence of excitatory intratelencephalic (IT) neurons involved in ASD is that the receptor tyrosine kinase MET, an autism risk allele, is expressed in IT neurons, and carriers show atypical activation or deactivation responses to social stimuli (Shepherd, 2013). Subtype-2 is uniquely enriched in astrocyte, Glutamatergic_L6b, microglia and oligodendrocyte progenitor cells (OPC) for subtype-2. Disruption of astrocyte function in ASD may affect proper neurotransmitter metabolism, synaptogenesis, and the state of brain inflammation (Vakilzadeh & Martinez-Cerdeño, 2023). Neuropathological studies of ASD post-mortem tissues have shown evidence of increased microglial activation and neuroinflammation in multiple brain regions (Fan et al., 2023). Mature oligodendrocytes originate from OPCs, and the aberrant development of white matter found in patients with ASD and disrupted protein levels in oligodendrocytes in ASD mice models (Galvez-Contreras et al., 2020). These results suggest distinct genetic signatures among the three subtypes.

The current study has several limitations. Firstly, we did not include a replication cohort due to the limited number and quality of neuroimaging data. We instead used bootstrapping and calculated the similarity matrix to maximize the stability, which we believe is a good alternative that addresses generalizability. Secondly, this study used linear mixed-effects models to account for the age effect, while further exploration of potential nonlinear effects is important. Third, the dataset (ABIDE) contained limited data on behavior/symptoms. These brain structural network-based subtypes may differ in other behavioral dimensions not captured by the ADOS and ADIR. Finally, the dataset that we used did not have the clinical intervention information. It could be more informative if differences in the symptoms or cognitive ability improvement could be investigated among subtypes after a period of intervention.

In conclusion, by using a data-driven approach on a relatively large cohort of individuals with ASD, we found three MRI subtypes characterized by different patterns of structural network as measured by MSN, behavioral, and functional connectivity profiles. These findings could be potentially helpful for advancing personalized medicine or other intervention approaches in ASD.

## References

[1] Abbott, A. E. (2016). Patterns of atypical functional connectivity and behavioral links in autism differ between default, salience, and executive networks. Cerebral Cortex, 26(10), 4034–4045.

[2] Arnatkevic iūtė, A. (2019). A practical guide to linking brain-wide gene expression and neuroimaging data. NeuroImage, 189, 353–367.

[3] Badre, D., & D’esposito, M. (2009). Is the rostro-caudal axis of the frontal lobe hierarchical? Nature Reviews Neuroscience, 10(9), 659–669.

[4] Bernhardt, B. C. (2016). Neuroimaging-based phenotyping of the autism spectrum. Social Behavior from Rodents to Humans, 341–355.

[5] Bigler, E. D. (2007). Superior temporal gyrus, language function, and autism. Developmental neuropsychology, 31(2), 217–238.

[6] Chandran, V. A. (2021). Brain structural correlates of autistic traits across the diagnostic divide: a grey matter and white matter microstructure study. NeuroImage: Clinical, 32, 102897.

[7] Chen, H. (2019). Parsing brain structural heterogeneity in males with autism spectrum disorder reveals distinct clinical subtypes. Human Brain Mapping, 40(2), 628–637.

[8] Chen, R. (2011). Structural MRI in autism spectrum disorder. Pediatric Research, 69(8), 63–68.

[9] Chen, Y.-Y. (2022). Excessive functional coupling with less variability between salience and default mode networks in autism spectrum disorder. Biological Psychiatry: Cognitive Neuroscience and Neuroimaging, 7(9), 876–884.

[10] Dale, A. M. (1999). Cortical surface-based analysis: I. Segmentation and surface reconstruction. NeuroImage, 9(2), 179–194.

[11] Delmonte, S. (2013). Functional and structural connectivity of frontostriatal circuitry in Autism Spectrum Disorder. Frontiers in Human Neuroscience, 7, 430.

[12] Di Martino, A. (2014). The autism brain imaging data exchange: towards a large-scale evaluation of the intrinsic brain architecture in autism. Molecular Psychiatry, 19(6), 659–667.

[13] Di Martino, A. (2017). Enhancing studies of the connectome in autism using the autism brain imaging data exchange II. Scientific Data, 4(1), 1–15.

[14] Fan, G. (2023). Microglia Modulate Neurodevelopment in Autism Spectrum Disorder and Schizophrenia. International journal of molecular sciences, 24(24), 17297.

[15] Fan, J. (2017). Spontaneous neural activity in the right superior temporal gyrus and left middle temporal gyrus is associated with insight level in obsessive-compulsive disorder. Journal of Affective Disorders, 207, 203–211.

[16] Filice, F. (2016). Reduction in parvalbumin expression not loss of the parvalbumin-expressing GABA interneuron subpopulation in genetic parvalbumin and shank mouse models of autism. Molecular Brain, 9, 1–17.

[17] Fortin, J.-P. (2018). Harmonization of cortical thickness measurements across scanners and sites. NeuroImage, 167, 104–120.

[18] Galvez-Contreras, A. Y. (2020). Role of oligodendrocytes and myelin in the pathophysiology of autism spectrum disorder. Brain sciences, 10(12), 951.

[19] Harika-Germaneau, G. (2022). Baseline clinical and neuroimaging biomarkers of treatment response to high-frequency rTMS over the left DLPFC for resistant depression. Frontiers in psychiatry, 13, 894473.

[20] Hawrylycz, M. J. (2012). An anatomically comprehensive atlas of the adult human brain transcriptome. Nature, 489(7416), 391–399.

[21] Hong, S.-J. (2018). Multidimensional neuroanatomical subtyping of autism spectrum disorder. Cerebral Cortex, 28(10), 3578–3588.

[22] Hyde, K. L. (2010). Neuroanatomical differences in brain areas implicated in perceptual and other core features of autism revealed by cortical thickness analysis and voxel-based morphometry. Human Brain Mapping, 31(4), 556–566.

[23] Jao Keehn, R. J. (2017). Impaired downregulation of visual cortex during auditory processing is associated with autism symptomatology in children and adolescents with autism spectrum disorder. Autism Research, 10(1), 130–143.

[24] Jiang, X. (2023). A brain structural connectivity biomarker for autism spectrum disorder diagnosis in early childhood. Psychoradiology, 3, kkad005.

[25] Jung, M. (2019). Decreased structural connectivity and resting-state brain activity in the lateral occipital cortex is associated with social communication deficits in boys with autism spectrum disorder. NeuroImage, 190, 205–212.

[26] Kana, R. K. (2006). Sentence comprehension in autism: thinking in pictures with decreased functional connectivity. Brain, 129(9), 2484–2493.

[27] Khundrakpam, B. S. (2017). Cortical thickness abnormalities in autism spectrum disorders through late childhood, adolescence, and adulthood: a large-scale MRI study. Cerebral Cortex, 27(3), 1721–1731.

[28] Koebis, M. (2019). LAMP5 in presynaptic inhibitory terminals in the hindbrain and spinal cord: a role in startle response and auditory processing. Molecular Brain, 12, 1–13.

[29] Lawrence, Y. (2010). Parvalbumin -, calbindin -, and calretinin - immunoreactive hippocampal interneuron density in autism. Acta Neurologica Scandinavica, 121(2), 99–108.

[30] Lepping, R. J. (2022). Visuomotor brain network activation and functional connectivity among individuals with autism spectrum disorder. Human Brain Mapping, 43(2), 844–859.

[31] Li, J. (2021). Cortical structural differences in major depressive disorder correlate with cell type-specific transcriptional signatures. Nature Communications, 12(1), 1–14.

[32] Li, W. (2017). Construction of individual morphological brain networks with multiple morphometric features. Frontiers in neuroanatomy, 11, 34.

[33] Liu, G. (2022). Two neuroanatomical subtypes of males with autism spectrum disorder revealed using semi-supervised machine learning. Molecular autism, 13(1), 1–14.

[34] London, E. B. (2014). Categorical diagnosis: a fatal flaw for autism research? Trends in Neurosciences, 37(12), 683–686.

[35] MacQueen, J. (1967). Some methods for classification and analysis of multivariate observations. Proceedings of the fifth Berkeley symposium on mathematical statistics and probability,

[36] Markello, R. D. (2021). Standardizing workflows in imaging transcriptomics with the abagen toolbox. eLife, 10, e72129.

[37] Markello, R. D. (2022). Neuromaps: structural and functional interpretation of brain maps. Nature methods, 19(11), 1472–1479.

[38] Monereo-Sánchez, J. (2021). Quality control strategies for brain MRI segmentation and parcellation: Practical approaches and recommendations-insights from the Maastricht study. NeuroImage, 237, 118174.

[39] Morgan, S. E. (2019). Cortical patterning of abnormal morphometric similarity in psychosis is associated with brain expression of schizophrenia-related genes. Proceedings of the National Academy of Sciences, 116(19), 9604–9609.

[40] Padmanabhan, A. (2017). The default mode network in autism. Biological Psychiatry: Cognitive Neuroscience and Neuroimaging, 2(6), 476–486.

[41] Palejwala, A. H. (2020). Anatomy and white matter connections of the lateral occipital cortex. Surgical and Radiologic Anatomy, 42, 315–328.

[42] Petrides, M. (2023). On the evolution of polysensory superior temporal sulcus and middle temporal gyrus: a key component of the semantic system in the human brain. Journal of Comparative Neurology, 531(18), 1987–1995.

[43] Pua, E. P. K. (2017). Autism spectrum disorders: Neuroimaging findings from systematic reviews. Research in Autism Spectrum Disorders, 34, 28–33.

[44] Rodríguez-Cruces, R. (2020). Multidimensional associations between cognition and connectome organization in temporal lobe epilepsy. NeuroImage, 213, 116706.

[45] Romero-Garcia, R. (2012). Effects of network resolution on topological properties of human neocortex. NeuroImage, 59(4), 3522–3532.

[46] Rosen, A. F. (2018). Quantitative assessment of structural image quality. NeuroImage, 169, 407–418.

[47] Seidlitz, J. (2018). Morphometric similarity networks detect microscale cortical organization and predict inter-individual cognitive variation. Neuron, 97(1), 231-247. e237.

[48] Shepherd, G. M. (2013). Corticostriatal connectivity and its role in disease. Nature Reviews Neuroscience, 14(4), 278–291.

[49] Skene, N. G., & Grant, S. G. (2016). Identification of vulnerable cell types in major brain disorders using single cell transcriptomes and expression weighted cell type enrichment. Frontiers in neuroscience, 10, 16.

[50] Stevenson, R. A. (2016). Keeping time in the brain: Autism spectrum disorder and audiovisual temporal processing. Autism Research, 9(7), 720–738.

[51] Tiveron, M.-C. (2016). LAMP5 fine-tunes GABAergic synaptic transmission in defined circuits of the mouse brain. PloS one, 11(6), e0157052.

[52] Vakilzadeh, G., & Martinez-Cerdeño, V. (2023). Pathology and astrocytes in autism. Neuropsychiatric disease and treatment, 841–850.

[53] Wang, Z. (2019). Resting-state brain network dysfunctions associated with visuomotor impairments in autism spectrum disorder. Frontiers in integrative neuroscience, 13, 17.

[54] Whitfield-Gabrieli, S., & Nieto-Castanon, A. (2012). Conn: a functional connectivity toolbox for correlated and anticorrelated brain networks. Brain connectivity, 2(3), 125–141.

[55] Wold, S. (1987). Principal component analysis. Chemometrics and intelligent laboratory systems, 2(1-3), 37–52.

[56] Xu, J. (2020). Specific functional connectivity patterns of middle temporal gyrus subregions in children and adults with autism spectrum disorder. Autism Research, 13(3), 410–422.

[57] Yeo, B. T. (2011). The organization of the human cerebral cortex estimated by intrinsic functional connectivity. Journal of neurophysiology.

[58] Zabihi, M. (2020). Fractionating autism based on neuroanatomical normative modeling. Translational Psychiatry, 10(1), 1–10.

[59] Zhou, Y. (2019). Metascape provides a biologist-oriented resource for the analysis of systems-level datasets. Nature Communications, 10(1), 1–10.

